# Tryptophanase disruption underlies the evolution of insect-bacterium mutualism

**DOI:** 10.1101/2024.08.16.608268

**Authors:** Yayun Wang, Minoru Moriyama, Ryuichi Koga, Kohei Oguchi, Takahiro Hosokawa, Hiroki Takai, Shuji Shigenobu, Naruo Nikoh, Takema Fukatsu

## Abstract

Animal-microbe symbioses are omnipresent, where both partners often gain benefits as mutualists. How such mutualism has evolved between originally unrelated organisms is of interest. Here we report that, using an experimental symbiotic system between the stinkbug *Plautia stali* and the model bacterium *Escherichia coli*, disruption of a single bacterial gene *tnaA* encoding tryptophanase makes *E. coli* mutualistic to *P. stali*. Survey of natural bacterial mutualists across wild populations of *P. stali* and other stinkbug species uncovered that their *Pantoea*-allied symbionts consistently lack *tnaA* gene. Some *Pantoea* species like *P. ananatis* retain *tnaA* gene and cannot establish symbiosis with *P. stali*, but *tnaA*-disrupted *P. ananatis* partially restored the symbiotic capability. When a natural *Pantoea* mutualist of *P. stali* was transformed with a functional *tna* operon, its symbiotic capability reduced significantly. Our finding suggests that tryptophanase disruption may have facilitated the evolution of gut bacterial mutualists in insects.

## Introduction

Microbial symbioses prevail the biological kingdoms, in which the diverse relationships encompass parasitism, commensalism and mutualism (1,2). Among them, the most intimate symbioses are found among mutualistic ones, where the host and the symbiont constitute an integrated biological entity and suffer disadvantage without the partnership (3,4). Originally, such microbial symbionts must have had no relationship to their host organisms, plausibly having existed as environmental microbes. It is of fundamental evolutionary interest how ordinary free-living microbes have become indispensable mutualists. How many and what mutations are required for the evolution of mutualism? How quickly does the evolution of mutualism proceed? To address these questions, experimental evolutionary approaches may provide valuable insights (5–12).

Recently, an experimental evolutionary system consisting of an insect *Plautia stali* as host and a bacterium *Escherichia coli* as symbiont was established, which brought about an unprecedented insight into an early stage of the evolution of mutualistic symbiosis (13,14). The stinkbug *P. stali* possesses a midgut symbiotic organ full of a specific bacterial symbiont of the genus *Pantoea*, which is essential for growth and survival of the host insect (15–18). The famous model bacterium *E. coli* is a component of mammalian gut microbiome and has no prior relationship to the insect (19,20). However, when symbiont-deprived newborn nymphs of *P. stali* were experimentally inoculated and maintained with a hyper-mutating *E. coli* strain, multiple evolutionary lines exhibited significantly improved adult emergence rate and body color within several months to a year, indicating rapid and recurrent evolution of “mutualistic” *E. coli* (13). Analysis of the independently-evolved mutualistic *E. coli* lines revealed that single loss-of-function mutations on *cyaA* and *crp* genes that convergently disrupt the carbon catabolite repression (CCR) global transcriptional regulator system, which is involved in bacterial metabolic switching in response to nutritional stresses (21,22), are responsible for the mutualistic host phenotypes, uncovering that elaborate mutualistic symbiosis can evolve very easily and rapidly by a single gene mutation (13).

The finding that only a single gene disruption is sufficient for making *E. coli* an insect mutualist is certainly striking, but it should be noted that disruption of the CCR pathway globally affects the expression levels of over 500 downstream genes encoded on the *E. coli* genome (23). Hence, we expected that the causative genes directly responsible for the improved host phenotypes may be identified among the downstream *E. coli* genes under the CCR regulation, although the nature of such genes was totally unknown. Here we report the identification of a bacterial enzyme gene under the CCR regulation whose disruption underpins the evolution of the insect-bacterium mutualism.

## Results

### No CCR disruption in natural symbionts of *P. stali*

In Japanese populations of *P. stali*, six *Pantoea*-allied symbiotic bacteria, *Pantoea* spp. A, B, C, D, E and F (abbreviated as Sym A and Sym B for uncultivable ones, and Sym C, Sym D, Sym E and Sym F for cultivable ones) are present (15). While all the symbiotic bacteria can support normal growth and reproduction of *P. stali*, genome sequencing revealed that their genomes consistently retain the intact CCR pathway genes *cyaA* and *crp* (Table S1). These observations strongly suggested that the CCR disruption, which was observed in the laboratory evolution of mutualism with hyper-mutating *E. coli* (13), is not involved in the evolution of mutualistic symbionts in natural populations of *P. stali*.

### Survey of downstream *E. coli* genes affected by CCR disruption

Considering that disruption of the CCR global transcriptional regulator system generally affects expression levels of hundreds of genes encoded on bacterial genomes (23), it seemed plausible that some genes downstream of the CCR pathway may actually be responsible for the mutualistic phenotypes of the CCR disruptive *E. coli* mutants. In this context, we focused on 55 CCR-regulated genes that were identified to be commonly down-regulated in two independent mutualistic *E. coli* evolutionary lines, CmL05 and GmL07, identified in our previous study (13). These genes consisted of diverse array of functional genes such as transporter genes for non-glucose sugars, carbohydrate metabolism genes, quorum sensing genes, extracellular matrix production genes, transcription factor genes and others (Fig. S1).

### Elevated tryptophan level after evolution of mutualism as well as CCR disruption in *E. coli*

On the other hand, in order to gain insight into metabolic aspects of the mutualistic evolutionary and mutant *E. coli* lines, we conducted comparative transcriptomic and metabolomic analyses of the *E. coli*-infected *P. stali*. A promising clue came from quantitative analysis of free amino acids in *P. stali* infected with the CCR disruptive *E. coli* mutants Δ*cyaA* and Δ*crp*, in which a specific essential amino acid, tryptophan, exhibited over ten times higher levels in hemolymph and symbiotic organ in comparison with the insects infected with the control *E. coli* strain Δ*intS* (Fig. 1a, b; Fig. S2). Of the 55 candidate genes, two genes were related to tryptophan: *tnaA* encoding tryptophanase and *tnaB* encoding a component of tryptophan transporter (Fig. S1). We obtained deletion mutants of these genes, Δ*tnaA* and Δ*tnaB*, and inoculated the *E. coli* mutants to *P. stali*. Then, tryptophan levels in hemolymph and symbiotic organ were significantly elevated in the Δ*tnaA*-infected insects but not in the Δ*tnaB*-infected insects (Fig. 1a, b; Fig. S2). Here it should be noted that the elevated tryptophan levels in the Δ*tnaA*-infected insects were comparable to those in the insects infected with the CCR disruptive mutants Δ*cyaA* and Δ*crp*, comparable to those in insects infected with the mutualistic evolutionary *E. coli* strain CmL05 (13), and also comparable to those in the normal symbiotic insects with the natural *Pantoea* symbiont Sym A (Fig. 1a, b; Fig. S2; Fig. S3). Tryptophanase assay confirmed that not only Δ*tnaA* but also Δ*cyaA*, Δ*crp* and CmL05 lost the tryptophanase activity whereas ΔintS and Δ*tnaB* were tryptophanase-positive (Fig. S4).

**Fig. 1.**
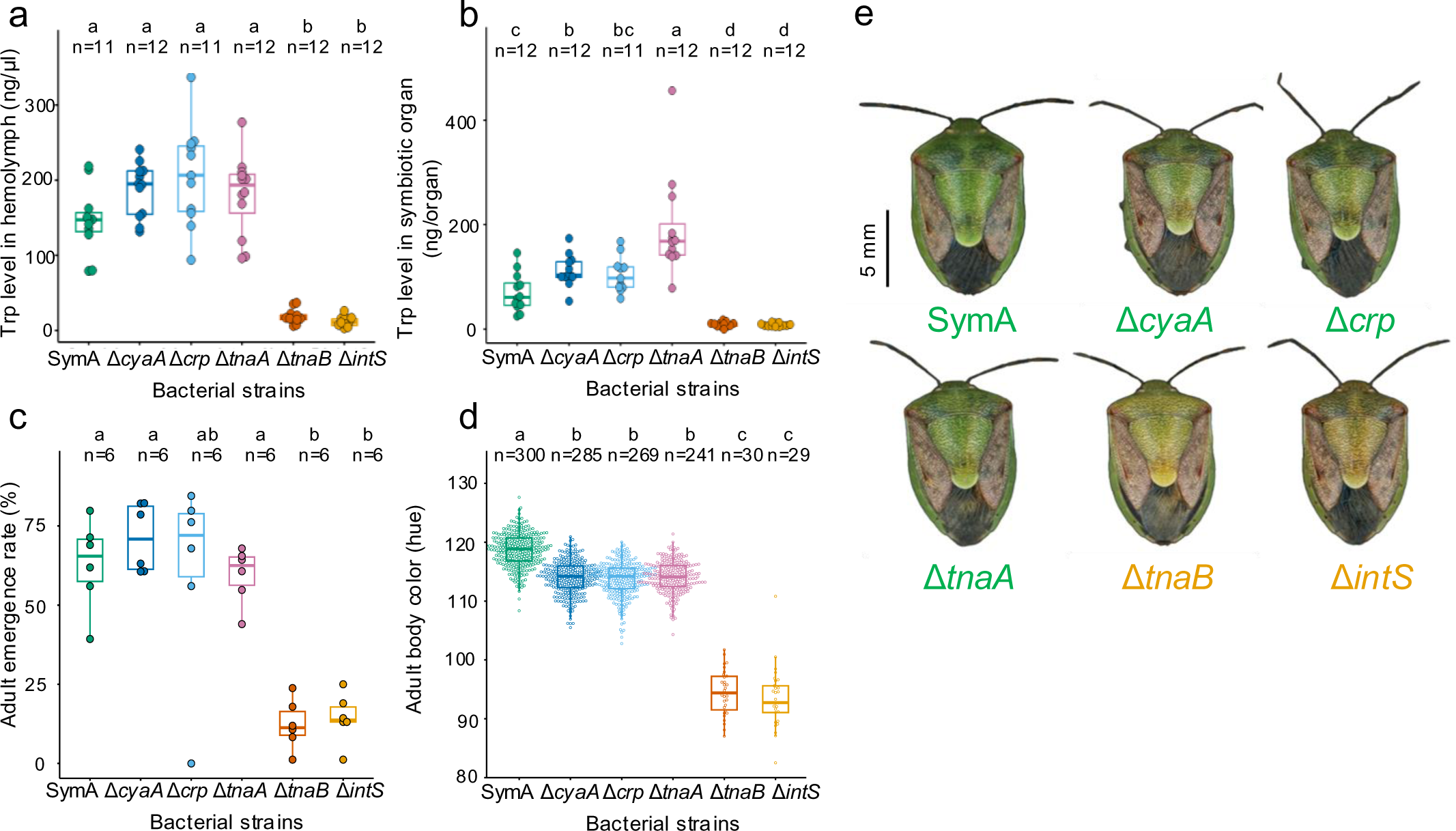
Improved phenotypes of *P. stali* infected with tryptophanase-disrupted Δ*tnaA* mutant of *E. coli*. (**a-d**) Effects of knockout mutants of *E. coli*, Δ*cyaA*, Δ*crp*, Δ*tnaA* and Δ*tnaB*, on tryptophan level in hemolymph (**a**), tryptophan level in symbiotic organ (**b**), adult emergence rate (**c**) and body color (**d**) when inoculated to *P. stali*. Note that Sym A, the natural symbiont of *P. stali* (15), comprises a mutualistic positive control, whereas Δ*intS E. coli* represents a non-beneficial negative control (13). Different alphabetical letters (a, b, c) indicate statistically significant differences (pairwise Wilcoxon rank-sum test with Hommel’s correction: *P* > 0.05). (**e**) External appearance of the adult insects infected with Sym A, Δ*cyaA*, Δ*crp*, Δ*tnaA*, Δ*tnaB* and Δ*intS* obtained in the study.

### Improved host phenotypes induced by tryptophanase disruption in *E. coli*

Furthermore, the Δ*tnaA*-infected insects exhibited significantly high adult emergence rates and remarkably greenish body color in comparison with the Δ*intS*-infected control insects, which were comparable to those of the Δ*cyaA-* and Δ*crp*-infected insects, and also comparable to those of the normal symbiotic insects with the natural *Pantoea* symbiont Sym A (Fig. 1c-e; Fig. S5a, b). On the other hand, the Δ*tnaA*-infected insects showed no significant improvement in body size in comparison with the Δ*intS*-infected control insects, which was also the case for the Δ*cyaA-* and Δ*crp*-infected insects (Fig. S5c, d). These results uncovered that (i) disruption of tryptophanase gene in *E. coli*, which is under the CCR regulation, significantly improves survival and body color of infected *P. stali*, (ii) the improved host performance due to infection with the CCR disruptive mutants, Δ*cyaA* and Δ*crp*, is largely attributable to down-regulation of the downstream *tnaA* gene, and (iii) tryptophanase disruption is a pivotal mechanism that underpins the evolution of *P. stali*-*E. coli* mutualism.

### On mechanism of improved host performance by tryptophanase disruption in *E. coli*

Figure 2 summarizes the results as to what molecular mechanisms are involved in the laboratory evolution of *P. stali*-*E. coli* mutualism. Before the evolution of mutualism, bacterial CCR operates, tryptophanase is expressed, tryptophan is broken down, and host suffers poor performance (Fig. 2a). After the evolution of mutualism, bacterial CCR is disrupted, tryptophanase is suppressed, tryptophan accumulates, and host shows good performance (Fig. 2b). Here, then, why does tryptophanase disruption in the symbiotic *E. coli* end up with the improved performance of the host *P. stali*? Considering that tryptophanase converts tryptophan into indole, pyruvate and ammonium (24), we conceived two plausible hypotheses, which are not necessarily mutually exclusive. One hypothesis is that tryptophanase disruption suppresses toxic indole production and thereby improves the host fitness, on the ground that perturbation of tryptophan metabolism mediated by gut microbiota toward the indole pathway tends to be linked to pathology and disease (25). Another hypothesis is that tryptophanase disruption results in accumulation of the essential amino acid tryptophan and thereby contributes to the host fitness, given that symbiont-mediated provisioning of tryptophan is important for diverse plant-sucking insects (26).

**Fig. 2.**
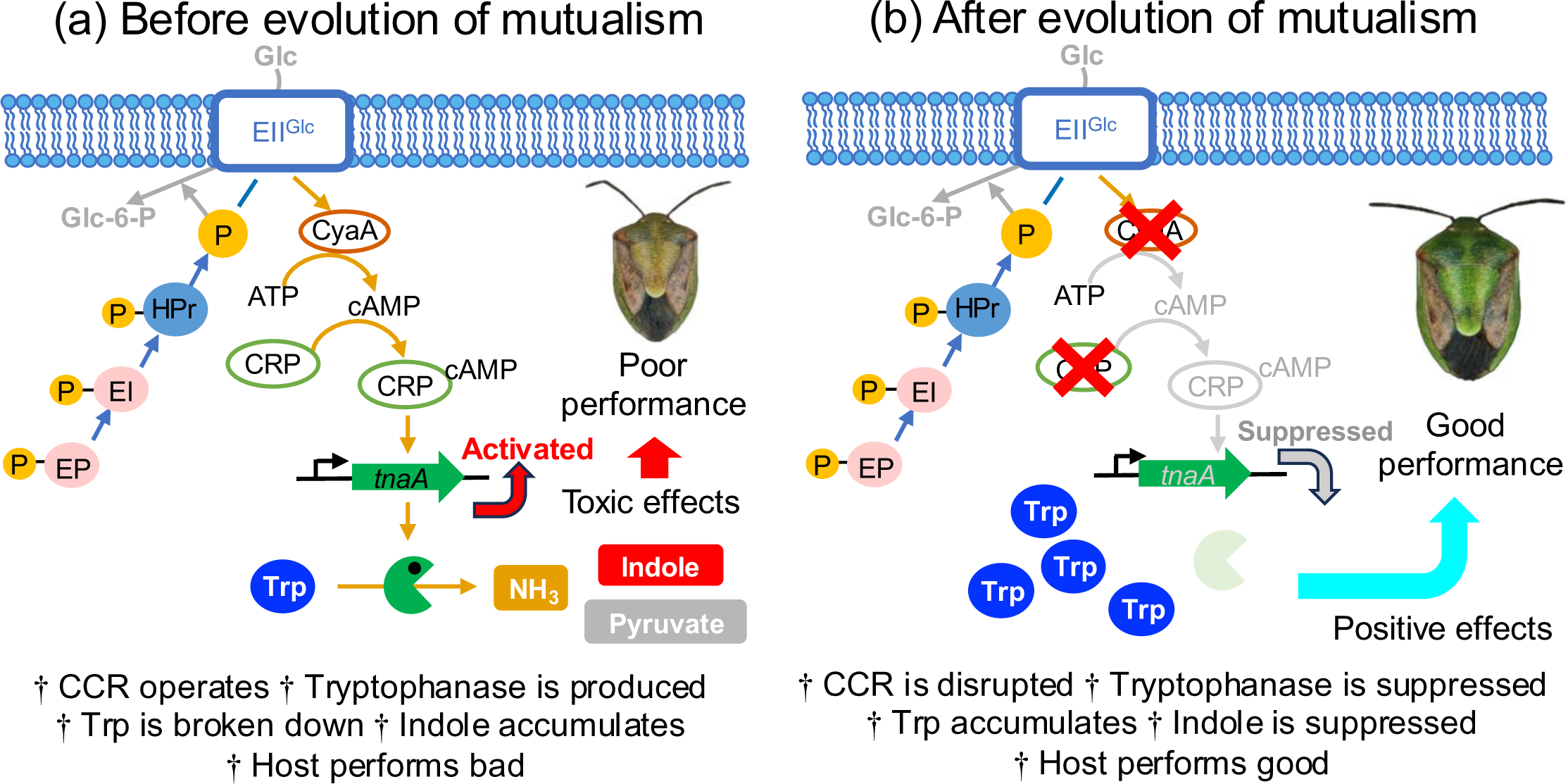
Molecular mechanisms underlying the evolution of *P. stali*-*E. coli* mutualism uncovered in this study. (**a**) Before the evolution of mutualism. (**b**) After the evolution of mutualism.

### Effects of indole and tryptophan feeding

For testing these hypotheses, we administrated different concentrations of indole and tryptophan to *P. stali* nymphs infected with the tryptophanase-deficient Δ*tnaA E. coli* and those infected with the control Δ*intS E. coli* via drinking water. As indole doses were elevated, adult emergence rates declined in both the Δ*tnaA*-infected insects and the Δ*intS-*infected insects, where the level of decline was less conspicuous in the Δ*tnaA*-infected insects than in the Δ*intS-* infected insects (Fig. 3a, b). These results favored the notion that indole accumulation is detrimental to growth and survival of *P. stali*. As tryptophan doses were elevated, adult emergence rates were not affected in the Δ*tnaA*-infected insects (Fig. 3c) but suppressed in the Δ*intS*-infected insects (Fig. 3d). These results seemed unexpected at a glance considering that tryptophan is an essential amino acid. However, it should be noted that the laboratory insects are provided with highly nutritious foods (raw peanuts), tryptophan feeding may thus lead to excessive tryptophan intake, and the tryptophanase-positive Δ*intS E. coli* may convert the excess tryptophan into toxic indole. Quantification of tryptophan and indole in hemolymph samples of these experimental insects revealed that (i) the Δ*tnaA*-infected insects exhibited little hemolymphal indole, (ii) by contrast, the Δ*intS*-infected insects showed significantly higher levels of hemolymphal indole, (iii) in the Δ*intS*-infected insects, indole/tryptophan feeding tended to result in elevated levels of hemolymphal indole, and (iv) in the Δ*intS*-infected insects, tryptophan levels were consistently low (Fig. 3e, f). These results accounted for the observations that not only indole feeding but also tryptophan feeding ended up with negative fitness consequences preferentially in the Δ*intS*-infected insects (Fig. 3a-d). Metabolomic analysis confirmed that the higher hemolymphal tryptophan levels and the lower hemolymphal indole levels were observed in the insects infected with the CCR-deficient mutant and evolutionary *E. coli* strains, Δ*cyaA* and CmL05G13, in comparison with the Δ*intS*-infected insects (Fig. S6a-c). It was also shown that some tryptophan-derived metabolites, such as 5-hydroxytryptamine, kynurenine, 3-hydroxykynurenine, indole-3-acetic acid and indole-3-carboxylic acid, tended to exhibit higher hemolymphal levels in the insects infected with the CCR-deficient mutant and evolutionary *E. coli* strains, Δ*cyaA* and CmL05G13, in comparison with the Δ*intS*-infected insects (Fig. S6d-k).

**Fig. 3.**
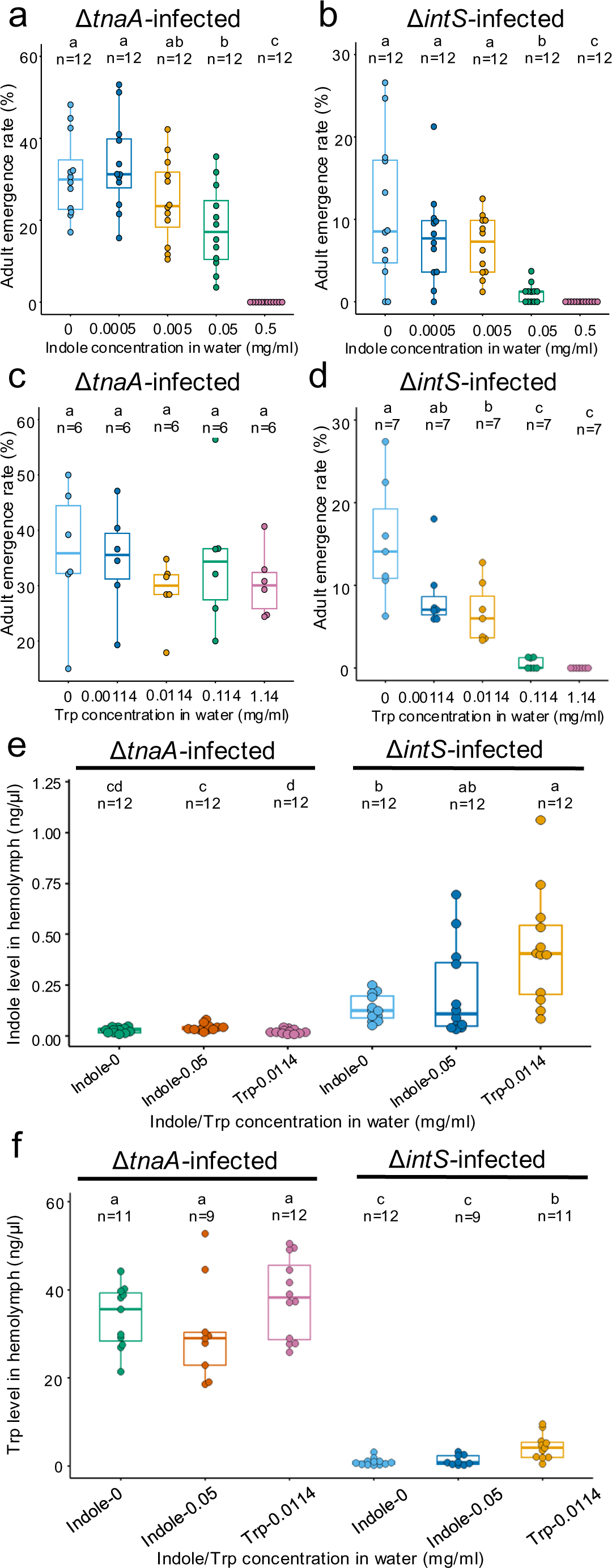
Effects of oral administration of indole and tryptophan on *P. stali* infected with tryptophanase-disrupted and control *E. coli*. (**a, b**) Effects of indole feeding via drinking water on growth and survival of *P. stali* infected with tryptophanase-disrupted Δ*tnaA E. coli* (**a**) and control Δ*intS E. coli* (**b**). (**c, d**) Effects of tryptophan feeding via drinking water on growth and survival of *P. stali* infected with tryptophanase-disrupted Δ*tnaA E. coli* (**c**) and control Δ*intS E. coli* (**d**). (**e, f**) Effects of indole/tryptophan feeding via drinking water on hemolymphal indole levels (**e**) and tryptophan levels (**f**) of *P. stali* infected with tryptophanase-disrupted Δ*tnaA E. coli*(**a**) and control Δ*intS E. coli*. Different alphabetical letters (a, b, c) indicate statistically significant differences (two-sided pairwise Wilcoxon rank-sum test with Hommel’s correction: *P* > 0.05).

### Absence of tryptophanase gene in natural symbiotic bacteria of *P. stali* and other stinkbugs

Given that tryptophanase disruption makes *E. coli* mutualistic to *P. stali* in laboratory, it is of interest whether natural symbiotic bacteria of stinkbugs retain *tnaA* gene or not. First, we inspected 6 genomes of natural *Pantoea*-allied symbionts Sym A, B, C, D, E and F of *P. stali*, in which no *tnaA* gene was found (Fig. 4; Table S1). Next, we determined 7 genomes of *Pantoea*-allied Sym C isolated from additional 7 Ryukyu island populations of *P. stali*, from which no *tnaA* gene was detected (Fig. 4; Table S1). Next, we determined 5 genomes of *Pantoea*-allied Sym C of other stinkbugs collected at Ryukyu islands, namely 3 local isolates from *Axiagastus rosmarus*, 1 isolate from *Lampromicra miyakona* and 1 isolate from *Scutellera amethystine,* all of which encoded no *tnaA* gene (Fig. 4; Table S1). Finally, using an inoculation and screening procedure with symbiont-free newborn nymphs of *P. stali* (15), we screened and isolated environmental bacteria capable of supporting growth of *P. stali* from soil samples collected at 5 Ryukyu islands, namely 6 isolates from Ishigaki Is., 2 isolates from Okinawa Is., 2 isolates from Miyako Is., 3 isolates from Yonaguni Is., and 4 isolates from Tokunoshima Is. All the environmental bacterial isolates potentially symbiotic to *P. stali* were phylogenetically placed in the genus *Pantoea* and devoid of *tnaA* gene in their genomes (Fig. 4; Table S1). Inoculation of these bacterial isolates to symbiont-free newborn nymphs of *P. stali* verified that they can support growth and survival of the host stinkbugs, where the stinkbug-derived isolates generally induced better host performance than the soil-derived isolates (Fig. S7). These results suggested that lack of *tnaA* gene may be related to the symbiotic capability of the *Pantoea*-allied bacteria to *P. stali*.

**Fig. 4.**
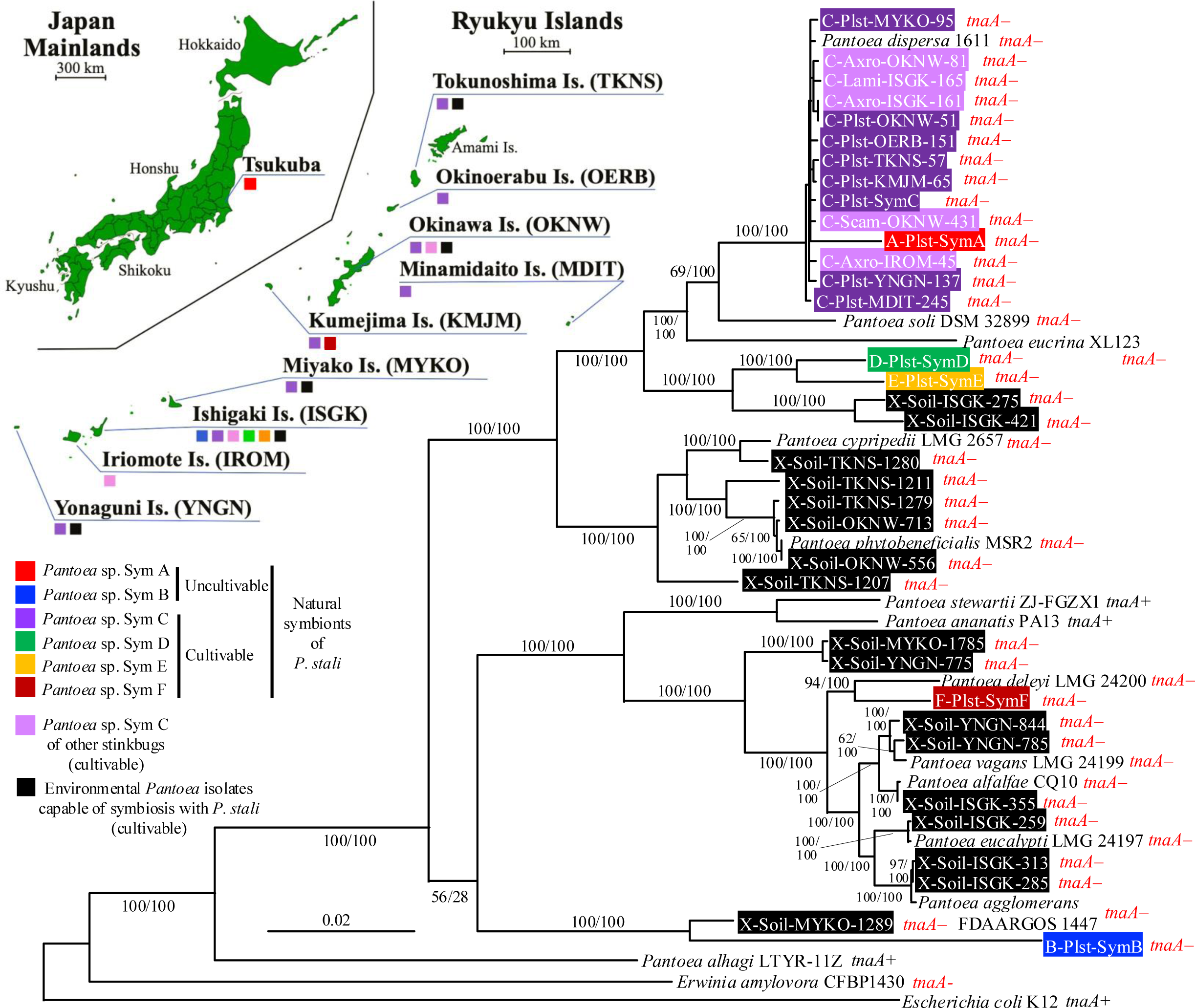
Molecular phylogenetic relationship of natural and potential mutualistic symbionts of *P. stali* and other stinkbugs. The maximum likelihood phylogeny is inferred from amino acid sequences of 106 concatenated essential single-core genes (35,647 aligned amino acid sites). Statistical support values for each clade are shown at the node in the order of maximum likelihood/Bayesian analyses. Collection localities are shown in the map of Mainland and Ryukyu islands of Japan. The colors for the bacterial taxon labels and the squares on the map correspond to the symbiont categories as depicted at bottom left (also see Table S1). Presence or absence of *tnaA* gene is shown beside the taxon labels.

### *Pantoea ananatis* with tryptophanase gene are incapable of symbiosis with *P. stali*

As of July 2024, 105 *Pantoea* genomes were deposited in the GenBank database. Among them, while 78 genomes lacked *tnaA* gene, 27 genomes retained *tnaA* gene, of which the majority, 19 genomes, were affiliated to *P. ananatis* (Table S2). We obtained 4 strains of *P. ananatis* from culture collections, which were all confirmed as *tnaA*-positive (Fig. S8a). When they were inoculated to symbiont-free newborn nymphs of *P. stali*, few adult insects emerged (Fig. S8b), indicating that the *tnaA*-carrying *P. ananatis* strains are incapable of establishing symbiosis with *P. stali*.

### Tryptophanase disruption improved symbiotic performance of *P. ananatis*

Is the inability of *P. ananatis* to establish symbiosis with *P. stali* relevant to tryptophanase gene on the bacterial genome? We generated a knockout mutant of *P. ananatis* JCM6986 by homologous recombination targeting *tnaA* gene (Fig. 5a). PCR detection confirmed deletion of *tnaA* gene (Fig. 5b) and enzymatic assay verified loss of tryptophanase activity in the mutant (Fig. 5c). When the Δ*tnaA* mutant of *P. ananatis* was inoculated to symbiont-free newborn nymphs of *P. stali*, nymphal survival and adult emergence rate significantly improved (Fig. 5d, e), although the adult emergence rate was only less than 10% on average (Fig. 5e). These results indicated that the non-symbiotic *Pantoea* strain becomes, though partially, mutualistic to *P. stali* by loss-of-function mutation of tryptophanase gene.

**Fig. 5.**
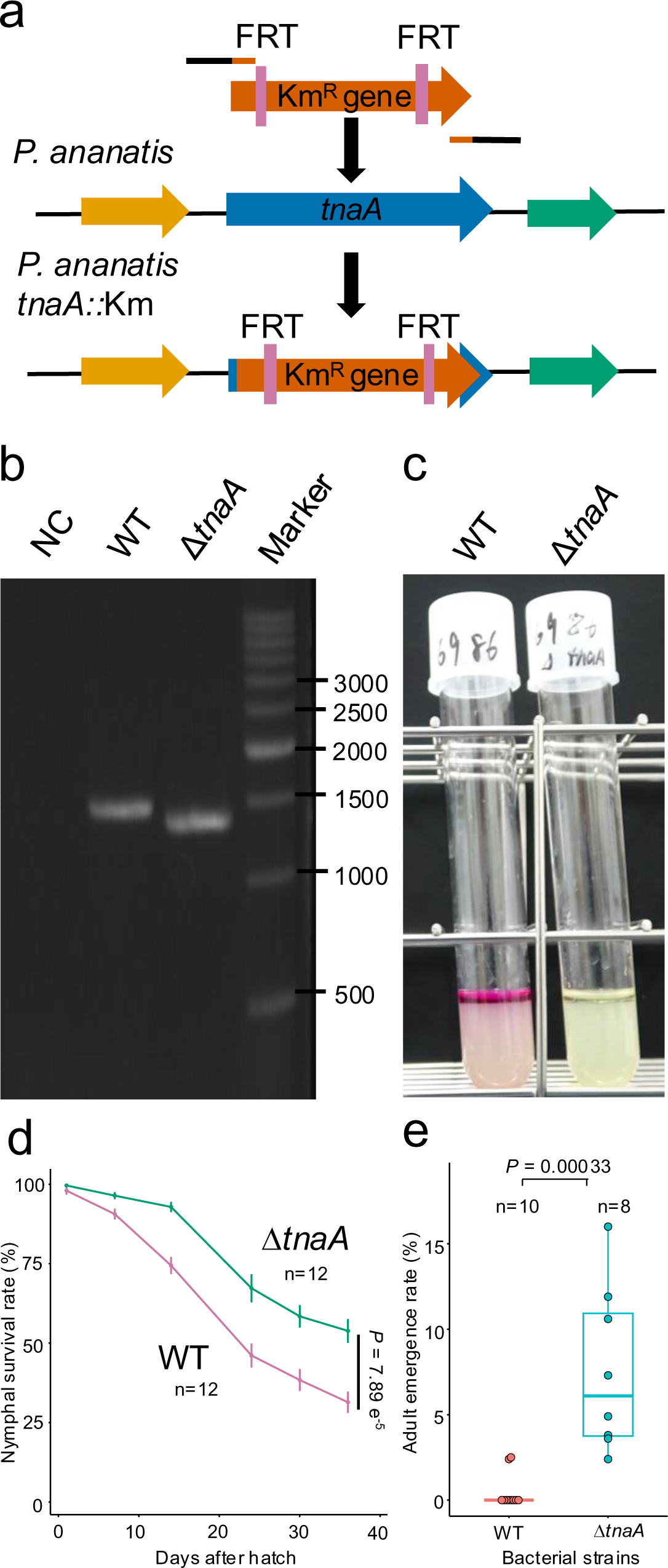
Knockout of tryptophanase gene *tnaA* in *P. ananatis* and effects on its symbiotic capability to *P. stali*. (**a**) Knockout scheme of *tnaA* gene by homologous recombination. (**b**) PCR check of homologous recombination. (**c**) Enzymatic assay of tryptophanase disruption. (**d**) Effects on survival curve. Statistical analysis was conducted on the 36^th^ day data by Welch’s two-sample test. (**e**) Effects on adult emergence rate. Statistical analysis was conducted by Student’s *t*-test.

### Natural *Pantoea* symbiont reduced symbiotic performance when transformed with functional tryptophanase

Finally, we artificially introduced a functional *tna* operon of *P. ananatis* into the genome of *Pantoea* sp. F (Sym F), a natural symbiont of *P. stali* that is cultivable and able to support host growth and survival (15), by homologous recombination targeting a presumably non-functional transposon-related gene *intB* (Fig. 6a; Fig. S9). The *intB*::*tnaAB* symbiont transformant exhibited a significant tryptophanase activity (Fig. 6b), verifying that the introduced *tna* operon is functioning in the transformed symbiont strain. When symbiont-free newborn nymphs of *P. stali* were inoculated with the tryptophanase-producing *intB*::*tnaAB* symbiont, their adult emergence rate and body color were negatively affected in comparison with those inoculated with the control Δ*intB* symbiont strain (Fig. 6c-f). In these insects, infection with the tryptophanase-producing *intB*::*tnaAB* symbiont caused higher tryptophan levels and lower indole levels than infection with the control Δ*intB* symbiont strain (Fig. 6g, h). These results indicated that the absence of tryptophanase gene may contribute to the mutualistic properties of the natural symbiont of *P. stali*.

**Fig. 6.**
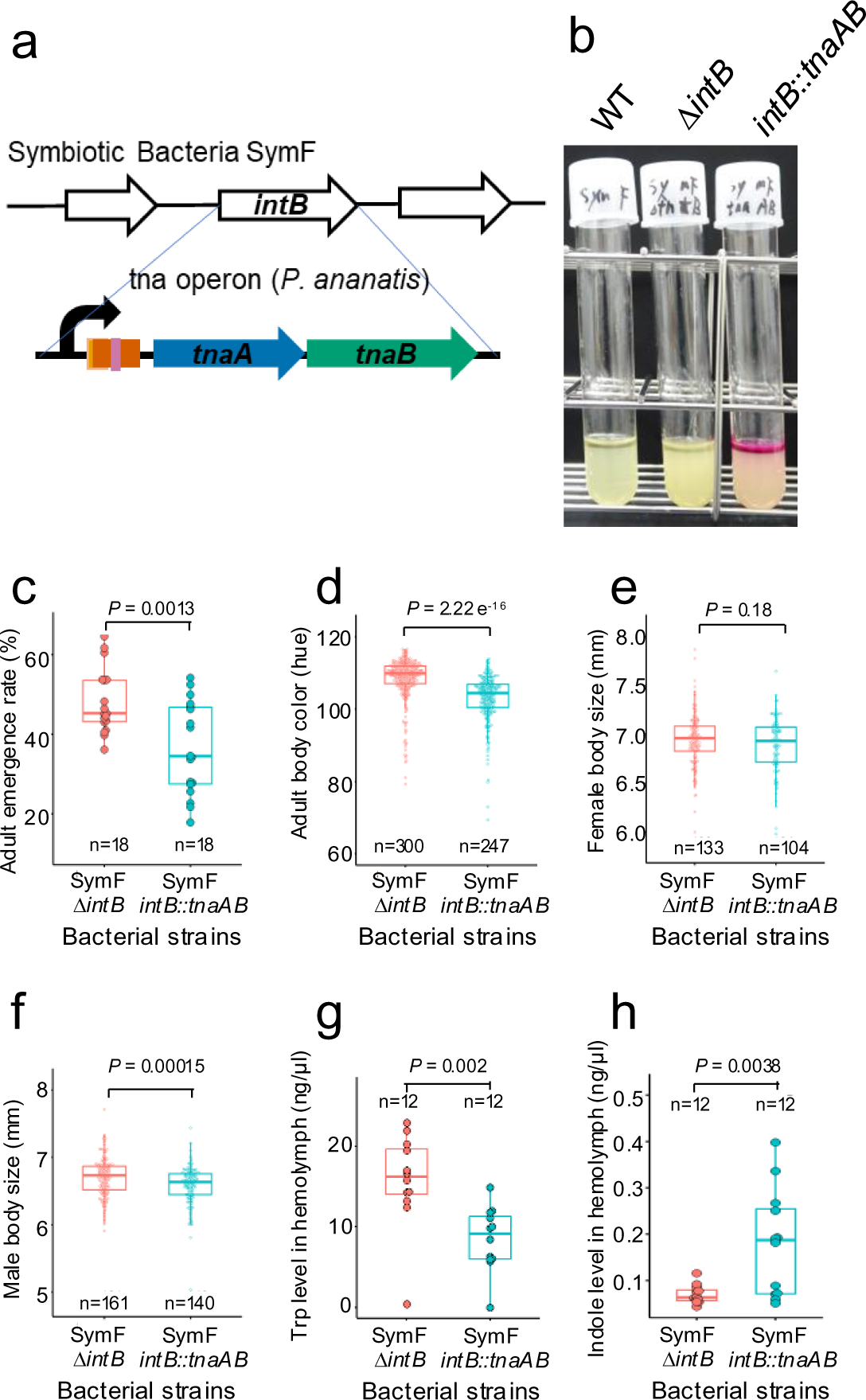
Transformation of natural symbiont *Pantoea* sp. F (Sym F) with functional *tna* operon and effects on its symbiotic capability to *P. stali*. (**a**) Introduction scheme of *tna* operon by homologous recombination. (**b**) Enzymatic assay of expression and functioning of introduced *tna* operon. (**c**) Effects on adult emergence rate. (**d**) Effects on adult body color. (**e**) Effects on adult female body size. (**f**) Effects on adult male body size. (**g**) Effects on tryptophan level in hemolymph. (**h**) Effects on indole level in hemolymph. Statistical analysis was conducted by Student’s *t*-test.

## DISCUSSION

Our previous study demonstrated that a single gene mutation, Δ*cyaA* or Δ*crp*, disrupting the bacterial CCR pathway makes *E. coli* mutualistic to *P. stali* (13), which led to the notion that elaborate mutualistic symbiosis can evolve more easily and rapidly than conventionally envisioned. On account of the diverse phenotypic changes observed with the mutualistic *E. coli* strains and mutants (13), we expected that multiple genes downstream of the CCR pathway may be involved in the evolution of *P. stali*-*E. coli* mutualism. Unexpectedly, however, we found that a single enzyme gene *tnaA* encoding tryptophanase, which is certainly under the CCR regulation, is the major effect gene whose disruption is sufficient for establishing *P. stali*-*E. coli* mutualism. This finding further corroborates the notion that elaborate mutualistic symbiosis can evolve easily and rapidly by a single gene mutation.

Plausibly, the tryptophanase disruption contributes to the host fitness via reduction of toxic indole, and also via accumulation of potentially limited essential amino acid tryptophan. It should be noted that diverse stinkbugs rely on their gut symbiotic bacteria for provisioning of essential amino acids and vitamins (27–30), and the *E. coli* genome encodes the genes needed for synthesis of all these nutrients (31). In plant-sucking aphids, the essential bacterial symbiont *Buchnera* encodes and amplifies synthetic genes for tryptophan on a plasmid (32, 33). Detailed physiological studies, for example those using a nutritionally defined artificial diet as developed for aphids (34, 35), are needed for further understanding of the insect-bacterium nutritional interactions and interdependency.

Our genomic and functional investigations uncovered that the loss of tryptophanase gene is not only underpinning the laboratory evolution of *P. stali*-*E. coli* mutualism but, plausibly, also involved in the evolution of bacterial mutualists of the genus *Pantoea* that have recurrently occurred in natural populations of *P. stali* and other stinkbugs (15, 36–38). Of course, a variety of genetic changes of both partners must have contributed to the establishment and maintenance of the stinkbug-bacterium mutualistic symbioses, only a part of which may be attributable to the loss of *tnaA* gene of the symbiont side. On account of the consistent absence of *tnaA* gene among the diverse stinkbug symbiont genomes, which encompass cultivable ones with large genome size to uncultivable ones with reduced genome size (Tables S1 and S3), we hypothesize that, although speculative, tryptophanase disruption may have facilitated the evolution of stinkbug-bacterium mutualism. Tryptophanase-deficient environmental *Pantoea* strains may have predisposed the establishment of symbiosis with stinkbugs. Alternatively, tryptophanase disruption may tend to occur at an early stage of the stinkbug-bacterium symbiosis in the course of symbiont genome degeneration. Anyway, it seems plausible that tryptophanase disruption acts as a pivotal mutation of the symbiont side that facilitates/canalizes/stabilizes the relationship toward mutualism.

By contrast, while loss-of-function mutations of *cyaA* and *crp*, which disrupt the CCR regulation, were identified as responsible for the evolution of *P. stali*-*E. coli* mutualism in laboratory (13), most of the *Pantoea*-allied natural symbiotic bacteria associated with *P. stali* and other stinkbugs, particularly those whose genomes are not so reduced, retain *cyaA* and *crp* genes in their genomes (Tables S1 and S3). These observations suggest that, in nature, disruption of the CCR pathway is generally not involved in the evolution of gut bacterial mutualists that are indispensable for the plant-sucking stinkbugs (39–41). Considering that the CCR regulation is important for bacterial adaptation to fluctuating environments by switching the main carbon source from depleted one to abundant one (21, 22), it is conceivable, although speculative, that the CCR disruption can evolve under stable environments like the laboratory condition, but it may be generally detrimental for such bacteria that are thriving under fluctuating natural environments. These observations provide an important lesson that the symbiotic evolution in laboratory does not necessarily reflect the symbiotic evolution in nature. Using the excellent model system for the experimental evolution of symbiosis between *P*. *stali* and *E. coli*, we have demonstrated that even a single bacterial gene mutation can facilitate the evolution of mutualism. In the real world, however, the processes and the mechanisms toward the evolution of mutualism must be much more complex entailing multiple genes and mutations of both host and symbiont sides. Integrative approaches to the evolution of mutualism as conducted in this study, in which experimental evolution in laboratory and natural diversity of symbiosis are jointly investigated, will lead to deeper understanding as to how elaborate mutualistic symbioses have been established and maintained.

## METHODS

### Insect samples, bacterial strains, and primers used in this study

Stinkbug samples and their symbiotic bacteria used in this study are listed (Table S1). Genome data of *Pantoea* isolates and stinkbug symbionts were retrieved from DNA databases (Tables S2 and S3). *P. ananatis* isolates JCM6986, JCM14682 and JCM15056 were obtained from the Japan Collection of Microorganisms, while the strain AJ13355 was provided by Ajinomoto Co., Inc. *E. coli* strains and mutants used in this study are listed (Table S4). The PCR primers used in this study are listed (Table S5).

### Insect rearing, symbiont sterilization and bacterial inoculation

For most experiments, a long-lasting laboratory strain of *P. stali* was used. The insects were reared in clean plastic or paper containers and fed with sterilized peanuts and sterilized water supplemented with 0.05% ascorbic acid in climate chambers at 25 ± 1°C under a long day regime of 16 h light and 8 h dark as described (42) (Fig. S10). For preparing symbiont-deprived newborn nymphs, collected egg masses were soaked in 4% formaldehyde for 20 min, kept twice in sterilized water for 10 min each, air-dried in a clean bench, and placed in sterile plastic Petri dishes with cotton balls. The Petri dishes were kept in an incubator at 25°C, where symbiont-free newborn nymphs emerged. Bacteria were cultured in liquid LB medium and diluted to OD600 = 0.1. The diluted culture medium (around 1.5 ml) was applied to cotton balls in each Petri dish, by which the symbiont-free newborn nymphs were allowed to orally acquire the bacterial suspension (Fig. S10a). After sucking bacteria-containing water, the first instar nymphs molted to second instar within 4-5 days without feeding, to which several pieces of sterilized peanuts and a 1.5 ml tube of sterilized water containing 0.05% ascorbic acid were introduced (Fig. S10b). Then, 3-4 days later, the mature second instar nymphs were transferred to a new rearing cage consisting of a paper container, a plastic lid with a large hole for ventilation, draining mesh for preventing insect escape, a 25 ml bottle of sterilized water containing 0.05% ascorbic acid, and sterilized peanuts (Fig. S10c, d). This rearing cage system, which was renewed every week, was devised for stably maintain the insect colonies in a good condition for an extended period. Six weeks after egg collection, the emerged adult insects were sexed, counted, kept in a refrigerator overnight, and image-scanned from their dorsal side using a scanner (EPSON GT-X980). Based on the scanned images, body color and body size of the insects were measured using an image analyzing software Natsumushi v.1.10 (43).

### Analysis of amino acids, tryptophan-derived metabolites, and indole

Each hemolymph sample was collected using a glass capillary (1 µl, Drummond) from the neck of an ice-anesthetized adult insect, suspended in 100 µL of 80% (v/v) methanol, and stored at ‒ 80°C until use. Each symbiotic midgut sample dissected from an adult insect was homogenized in 100 µl of 80% methanol and stored under the same condition. After adding homoarginine, homophenylalanine, [^15^N]-tryptophan, and 6-hydroxyindole as internal standards, each sample was centrifuged, and an aliquot of the supernatant was subjected to liquid chromatography and mass spectrometry (LC-MS) analysis of amino acids, tryptophan-derived metabolites, and indole. The detection and quantification of these metabolites were performed using an LC-MS system (Waters, ACQUITY UPLC H-class and Xevo G2-XS qTOF) with an electrospray ionization (ESI) source. Amino acid composition was measured after propyl-chloroformate derivatization as previously described (18, 30). For measurement of tryptophan and related compounds, the sample aliquots were concentrated under N2 flow, and resuspended in 0.05% (v/v) formic acid. Then, they were separated on a column (Waters, BEH C18, 1.7 µm, 2 mm x 100 mm) with a gradient elution of 0.05% formic acid and methanol. Each compound was selectively measured at a positive multiple reaction monitoring (MRM) mode. Since indole is not efficiently ionized under ESI, we derivatized it with p-dimethylaminocinnamaldehyde as described (44). The derivatized indole was separated on the same analytical column and quantified at a positive MRM mode.

### Feeding experiments with tryptophan and indole

L-tryptophan (FUJIFILM Wako Pure Chemical Corporation, Japan) and indole (Tokyo Chemical Industry Co., Ltd., Japan) were dissolved and serially diluted in sterilized water containing 0.05% ascorbic acid. As for tryptophan, concentrations of 1.14 x 10^0^, 10^-1^, 10^-2^ and 10^-3^ mg/ml were prepared. As for indole, concentrations of 0.5 x 10^0^, 10^-1^, 10^-2^ and 10^-3^ mg/ml were prepared. The experimental insects were reared with sterilized peanuts and the supplemented water as described (Fig. S10) except that the supplemented water was renewed every 3 or 4 days for minimizing deterioration of the supplemented reagents. Six weeks after egg collection, the emerged adult insects were counted and subjected to measurements of hemolymphal tryptophan and indole levels.

### Tryptophanase activity assay

A qualitative assessment of tryptophanase activity was conducted essentially as described (45). Each bacterial strain was cultured in 3 ml of LB liquid medium at 25°C with shaking at 180 rpm for 24 h. Then, 100 μl of KOVACS′ indole reagent (Sigma-Aldrich, St. Louis, MO) was added to the bacterial culture, by which produced indole was, if any, visualized by reddish color (see Fig. S4a). A quantitative assessment of tryptophanase activity during bacterial growth was conducted essentially as described (46). Each bacterial strain was cultured in LB liquid medium at 25°C with shaking at 180 rpm overnight, diluted with LB liquid medium to OD600 = 0.1, and dispensed to 24 test tubes as 1 ml aliquots. The test tubes were incubated at 25°C with shaking at 180 rpm, from which 3 samples were taken every hour and subjected to measurement of OD600. Then, the samples were centrifuged at room temperature at 10000 rpm for 3 min, and the supernatants were subjected to indole quantification using Indole Assay Kit (MAK326, Sigma-Aldrich, St. Louis, USA). (see Fig. S4b, c).

### Knockout of *tnaA* gene of *P. ananatis*

The presence of the *tnaA* gene in *P. ananatis* isolates was confirmed by PCR using the specific primers PA_tnaA_125F and PA_tnaA_1357R (Table S5). To knock out the *tnaA* gene in *P. ananatis* strain JCM6986, the primers PA_JCM6986_ΔtnaA_F and PA_JCM6986_ΔtnaA_R (Table S5) were used to amplify by PCR a kanamycin resistance gene (KmR) region from the *E. coli* Δ*intS* mutant. The PCR product was purified using QIAquick PCR Purification Kit (Qiagen, Netherlands). Transformation of *P. ananatis* JCM6986 with the plasmid pRed/ET (Gene Bridges GmbH, Germany) was performed using a Gene Pulser/MicroPulser (Bio-Rad Laboratories, USA). Subsequently, the *tnaA* gene in the *P. ananatis* genome was replaced by the KmR cassette (47). The successful insertion of KmR was confirmed by acquisition of kanamycin resistance in *P. ananatis*, and the loss of the pRed/ET plasmid was verified by the absence of tetracycline resistance. The deletion of the *tnaA* gene was further confirmed by specific PCR amplification using the primers PA_JCM6986_tnaA_check_F and PA_JCM6986_tnaA_check_R (Table S5). The resultant strain was named as *P. ananatis* JCM6986 Δ*tnaA* (Table S4).

### Transformation and expression of *tna* operon in *Pantoea* sp. F

Given that deletion of *tnaC* gene has been reported to result in constitutive expression of *tnaAB* genes (48), we first disrupted *tnaC* gene in *P. ananatis* JCM6986 by inserting a kanamycin resistance gene (KmR) region amplified by PCR using the primers PA_JCM6986_ΔtnaC_F and PA_JCM6986_ΔtnaC-RUT_R from the *E. coli* Δ*intS* mutant (Table S5) into the *tnaC* locus (Fig. S9). The successful deletion of *tnaC* was verified by specific PCR amplification with the primers PA_JCM6986_tnaC_check_F and PA_JCM6986_*tnaC*-RUT_check_R (Table S5). Next, the mutated *tna* operon from the resultant strain *P. ananatis* JCM6986 *tnaC*-RUT::Km (Fig. S9) was amplified using the primers SymF_intB/JCM6986_tnaAB_F and SymF_intB/JCM6986_tnaAB_R (Table S5). The *intB* gene of *Pantoea* sp. Plst-SymF (Table S1) was then replaced by the mutated *tna* operon from the PCR fragment, resulting in the construction of Plst-SymF *intB*::*tna* operon *tnaC*-RUT::Km (Fig. S9). Finally, the KmR cassette was excised using FLP recombinase expressed from the plasmid pFLP3. The loss of pFLP3 was facilitated by the *sacB*-based suicide gene system (49), thereby creating the final strain, Plst-SymF intB::tnaAB (Fig. S9; Table S4). The successful insertion of *tnaAB* was verified by specific PCR amplification with the primers SymF_intB_check_F and SymF_intB_check_R (Table S5). As a control strain in the infection experiments, the strain Plst-SymF Δ*intB* (Table S4) was constructed by the same protocol for the strain Plst-SymF *intB*::*tnaAB* except that the primers SymF_ΔintB_F and SymF_ΔintB_R (Table S5) were used to obtain a kanamycin resistance gene (KmR) region. The successful deletion of *intB* was verified by specific PCR amplification with the primers SymF_intB_check_F and SymF_intB_check_R (Table S5).

### Genome sequencing and analysis

DNAs of the uncultivable symbionts A-Plst-SymA and B-Plst-SymB (Table S1) were extracted from the symbiotic organs isolated from adult insects. DNA samples of the cultivable bacteria were extracted from overnight cultures in LB liquid medium at 25 °C. The extraction of DNA from the materials were conducted using DNeasy Blood and Tissue Kits (Qiagen, Netherland). The DNA samples were subjected to library preparation and sequencing using either PacBio RSII or PacBio Sequel sequencing systems (Pacific Biosciences of California, Inc., USA). As for the symbionts Sym A, C-Lami-ISGK-165 and C-Scam-OKNW-431 (Table S1), the DNA samples were additionally sequenced using MinION system in the combination with Ligation sequencing kit V14 and R10.4.1 flow cells (Oxford Nanopore Technologies, UK). Base calling of Nanopore long reads were conducted using Guppy ver. 6.5.7+ca6d6af (Oxford Nanopore Technologies) with minimap ver. 2.24-r1122 (50). *De novo* assembly was performed using Flye v.2.9.1 with default settings and an estimated genome size of 5.0 Mb (51). When libraries were sequenced with the PacBio RSII or Sequel system, raw PacBio reads were mapped to the draft assemblies via BLASR v.5.3.3 (52). The assembled sequences were then polished using the original PacBio reads with Arrow v.2.3.2 (53). The chromosomal genomes of A-Plst-SymA, C-Lami-ISGK-165, and C-Scam-OKNW-431 were not assembled into one circular contig due to the presence of long segmental repeats among different chromosomal regions. For these three strains, *de novo* assemblies of the ONT reads with Flye were used for gap-closing. The draft assemblies were polished with two rounds of medaka (v.1.4.1; https://github.com/nanoporetech/medaka). To reveal assembly errors, original sequencing reads were mapped to completing circular genomes using BWA v.0.7.17 (54). Possible sequence errors were identified using these mapped data via bam-readcont v.1.0.1 (55), and then manually inspected using IGV v.2.16.2 (56). Genome annotation was performed using DFAST v.1.2.20 (57).

### Molecular phylogenetic analysis

The genome sequences of the *Pantoea* symbionts, environmental *Pantoea* strains and allied bacteria were annotated using DFAST v.1.2.20. Then, the proteome sets were analyzed with publicly available ones using bcgTree v.1.2.0 (58), which automatically extracted 107 essential single-copy core genes from amino acid sequences of the whole genome data. Ambiguously aligned regions were trimmed by using Gblocks v.0.9.1b with manual inspection (59). In total, 106 gene alignments (one gene, *rpmH*, was excluded due to having missing data) were concatenated and a partitioning file was generated to mark the boundaries of each gene. Corresponding amino acid substitution models were estimated by the Bayesian information criteria using ModelTest-NG v.0.2.0 (60). The maximum likelihood analysis was conducted using RAxML-NG v0.9.0 (61). Bootstrap values were obtained with 1,000 resamples. The Bayesian inference analysis was conducted using MrBayes v.3.2.7a (62) with 10 million generations of Markov Chain Monte Carlo runs with sampling every 100 generations. The first 25 % samples were discarded as burn-in and the remaining trees were used to calculate posterior probabilities. The stationarity of the runs was assessed using Tracer v.1.7.2 (63).

### DATA AVAILABILITY

All genome sequencing data produced in this study were deposited in the DNA Data Bank of Japan (DDBJ) Sequence Read Archive (Table S1). The data have been deposited with links to BioProject accession no. PRJDB17425 in the DDBJ BioProject database. Source data are provided with this paper.

## ACKNOWLEDGEMENTS

We thank Dai Haraguchi and Yoshitomo Kikuchi for providing insect samples, and Tomoko Tanaka, Tomoko Matsushita, Tetsuhiro Hachikawa, Naomi Shimizu, Sakiko Toyoda, Mariko Taguchi and Megumi Kobayashi for technical assistance. This study was supported by the Japan Science and Technology Agency ERATO grant no. JPMJER1902 (T.F. and R.K), and the Japan Society for the Promotion of Science (JSPS) KAKENHI grant nos. JP25221107 (T.F. and R.K.), JP17H06388 (T.F. and S.S.), JP22128001 (T.F. and S.S.), and JP22128007 (T.F).

## AUTHOR CONTRIBUTIONS

Y.W., M.M., R.K. and T.F. conceived the project and designed the experiments. Y.W. conducted most of experimental works including bacterial infection, insect rearing, fitness evaluation, *E. coli* and *Pantoea* mutant generation, etc. M.M. performed metabolomic analyses of amino acids, indole and its derivatives, etc. R.K. conducted genomic and phylogenetic analyses of diverse stinkbug symbionts and soil-derived bacterial isolates. T.H. performed field survey to collect natural insects and soil samples, from which *Pantoea* strains symbiotic to *P. stali* were isolated.

K.O. and H.T. conducted inoculation experiments of insect-derived and soil-derived *Pantoea* isolates to *P. stali* and evaluated fitness consequences. S.S. and N.N. supported genomic and phylogenetic analyses. T.F. wrote the article with input from all authors. All authors approved the final version of the manuscript.

---

**Fig. S1.**
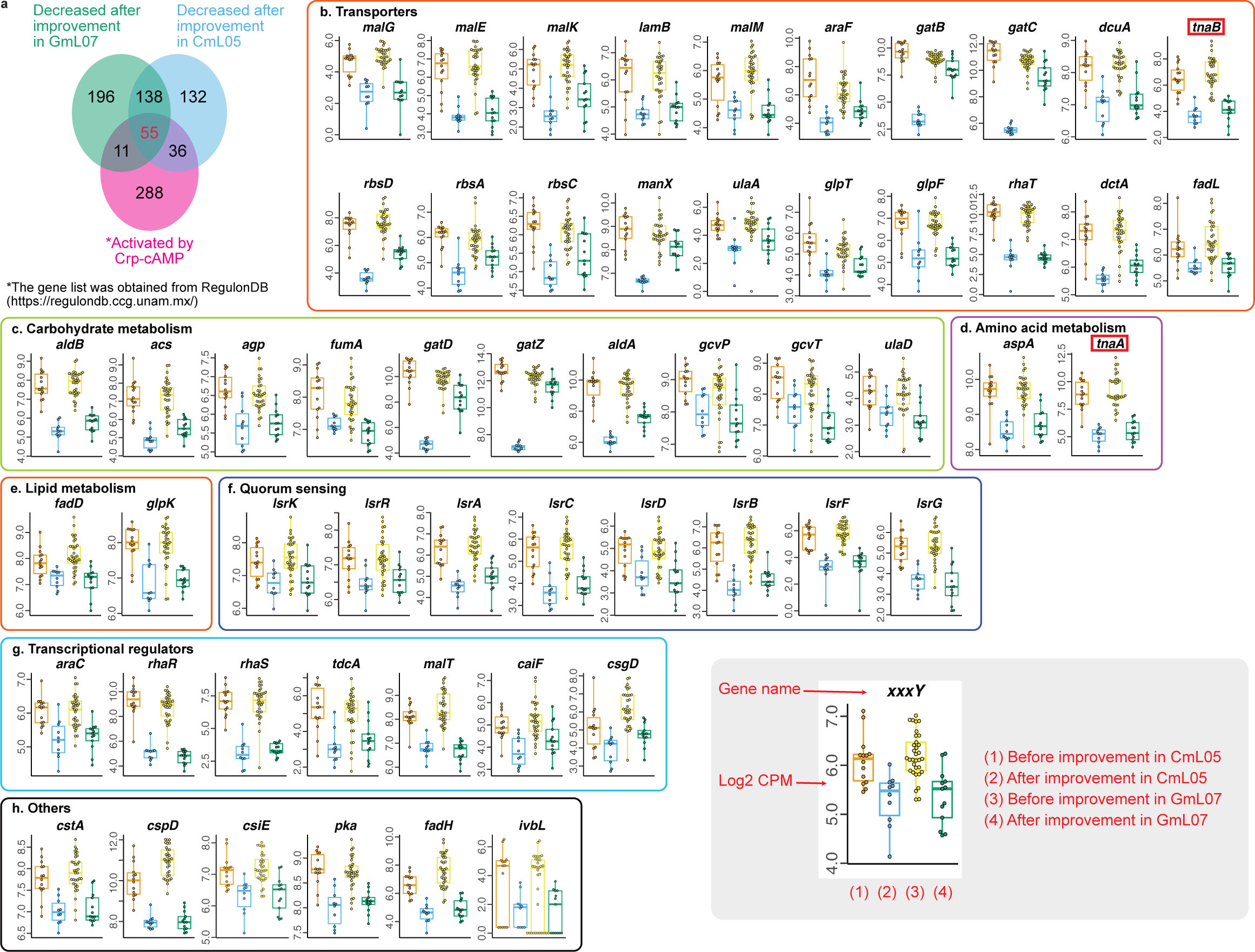
Genes commonly down-regulated in independent mutualistic *E. coli* evolutionary lines, CmL05 and GmL07, and also down-regulated by the CCR disruption in *E. coli*. (**a**) Venn diagram showing the numbers of commonly down-regulated genes. (**b-h**) Functional categories and expression levels of the commonly down-regulated genes. (**b**) Transporters. (**c**) Carbohydrate metabolism. (**d**) Amino acid metabolism. (**e**) Lipid metabolism. (**f**) Quorum sensing. (**g**) Transcriptional regulators. (**h**) Others. The inset figure at the bottom right represents the explanations of the elements in the plots. The numbers of the biological replicates and exact FDR *q*-values are provided with the source data.

**Fig. S2.**
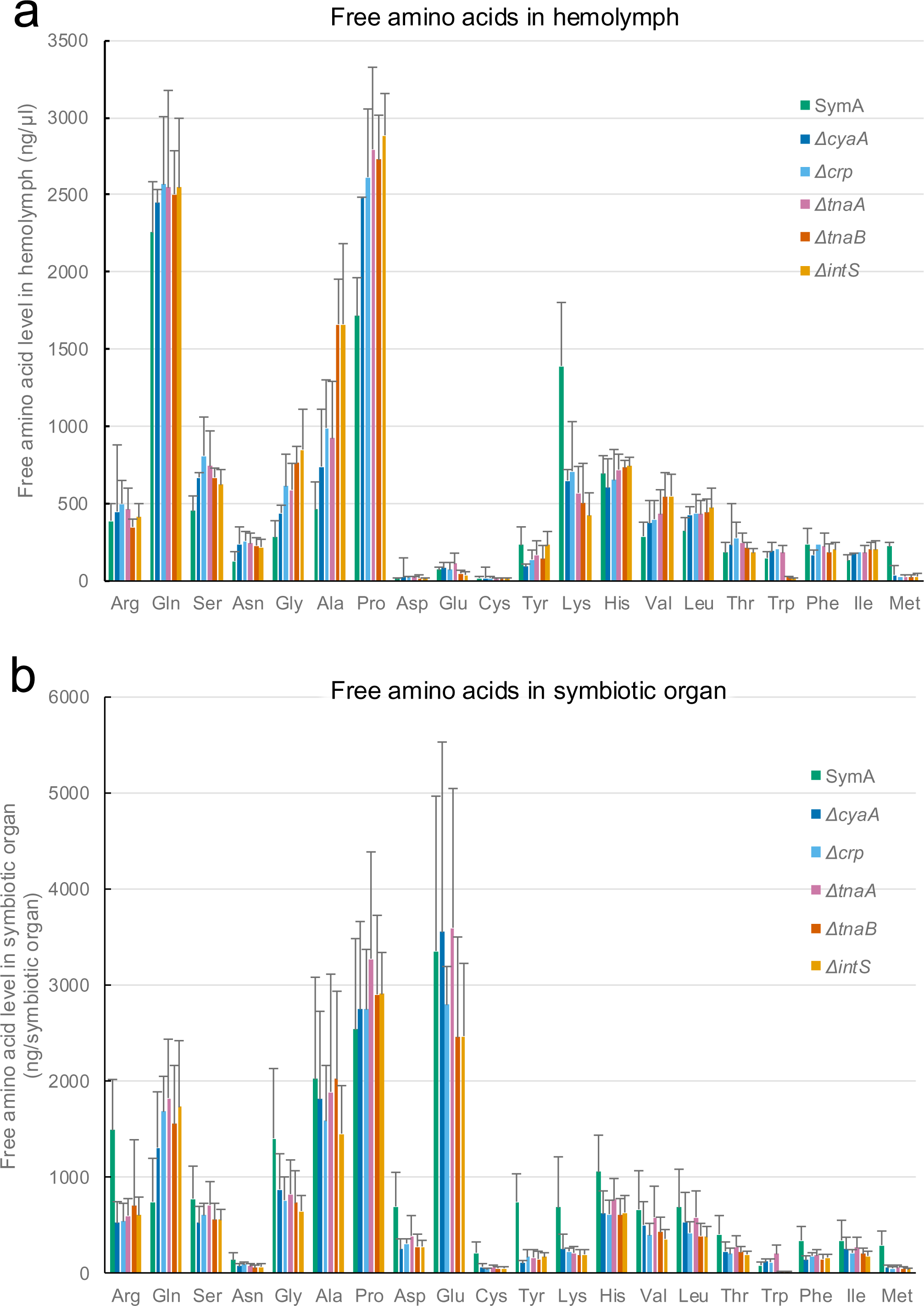
Levels of free amino acids in *P. stali* infected with a natural symbiont *Pantoea* sp. A (Sym A), mutant *E. coli* strains (Δ*cyaA*, Δ*crp*, Δ*tnaA* and Δ*tnaB*), and a wild-type *E. coli* strain (Δ*intS*). (**a**) Free amino acids in hemolymph. (**b**) Free amino acids in symbiotic organ. Note that tryptophan levels were drastically higher in the insects infected with Sym A, Δ*cyaA*, Δ*crp* and Δ*tnaA* than those infected with Δ*tnaB* and Δ*intS*.

**Fig. S3.**
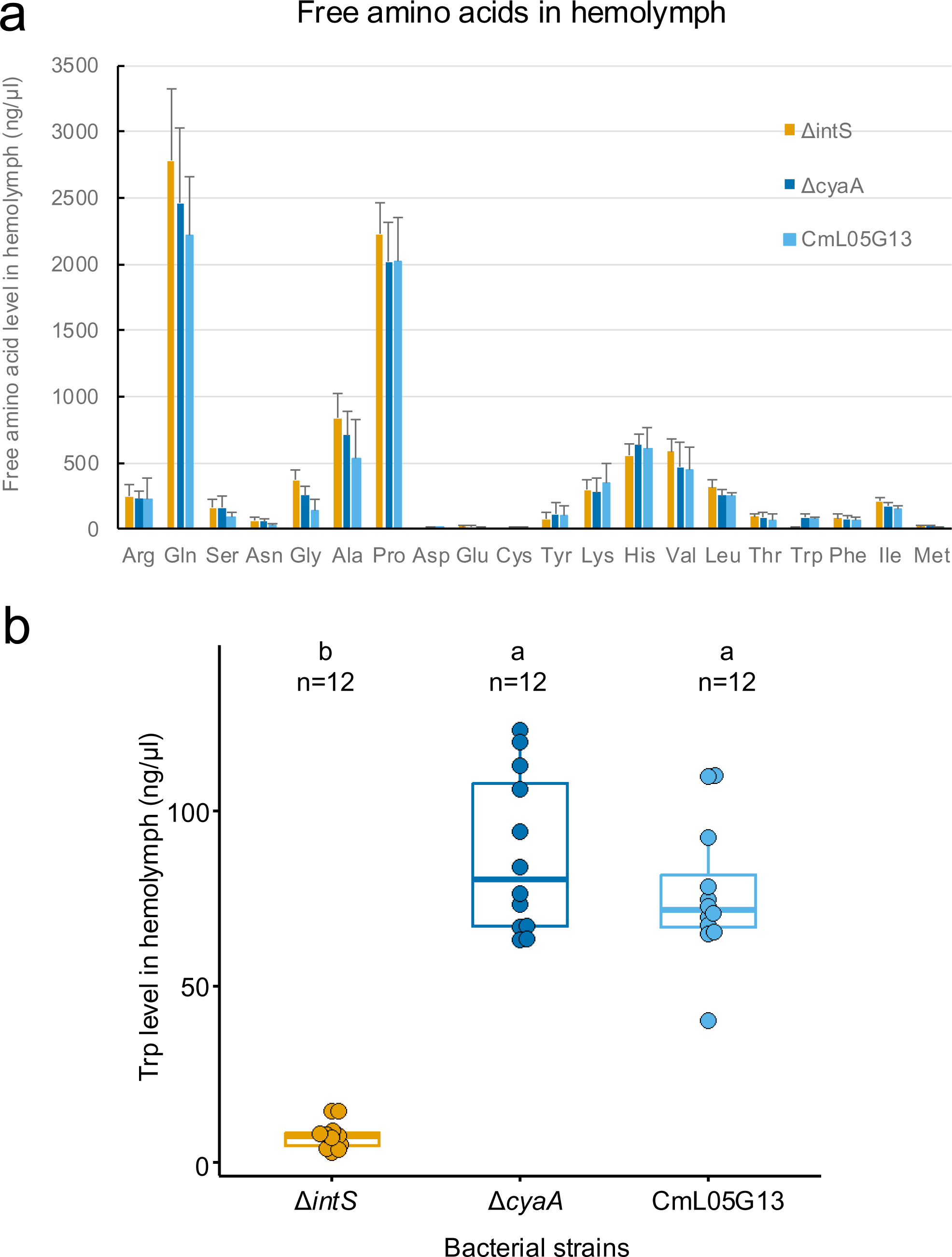
Levels of free amino acids in *P. stali* infected with a mutant *E. coli* strain (Δ*cyaA*), an evolutionary *E. coli* strain (CmL05G13), and a wild-type *E. coli* strain (Δ*intS*). (**a**) Free amino acids in hemolymph. (**b**) Comparison of tryptophan levels. Note that Δ*cyaA* and CmL05G13 are both CCR-deficient. Different alphabetical letters (a, b) indicate statistically significant differences (pairwise Wilcoxon rank-sum test with Hommel’s correction: *P* < 0.05).

**Fig. S4.**
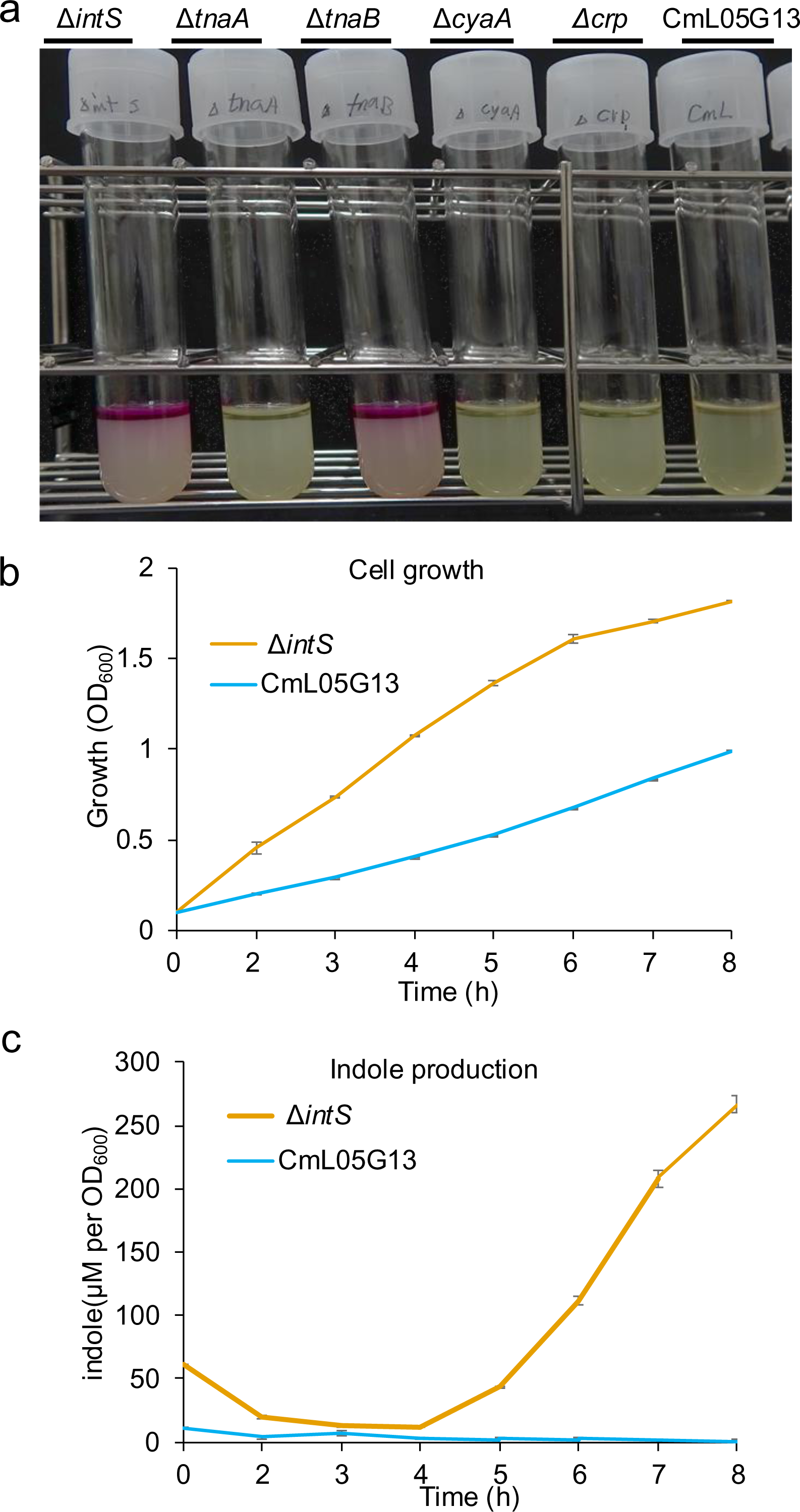
Tryptophanase assay of *E. coli* mutant strains. (**a**) Qualitative tryptophanase assay of the *E. coli* control strain Δ*intS*, the *E. coli* mutant strains Δ*tnaA*, Δ*tnaB*, Δ*cyaA* and Δ*crp*, and the evolutionary *E. coli* strain CmL05G13. Red color indicates tryptophanase activilty. (**b, c**) Quantitative cell growth dynamics (**b**) and indole production (**c**) of Δ*intS* and CmL05G13. Bacterial cell culture was conducted at 25°C in LB liquid medium with shaking.

**Fig. S5.**
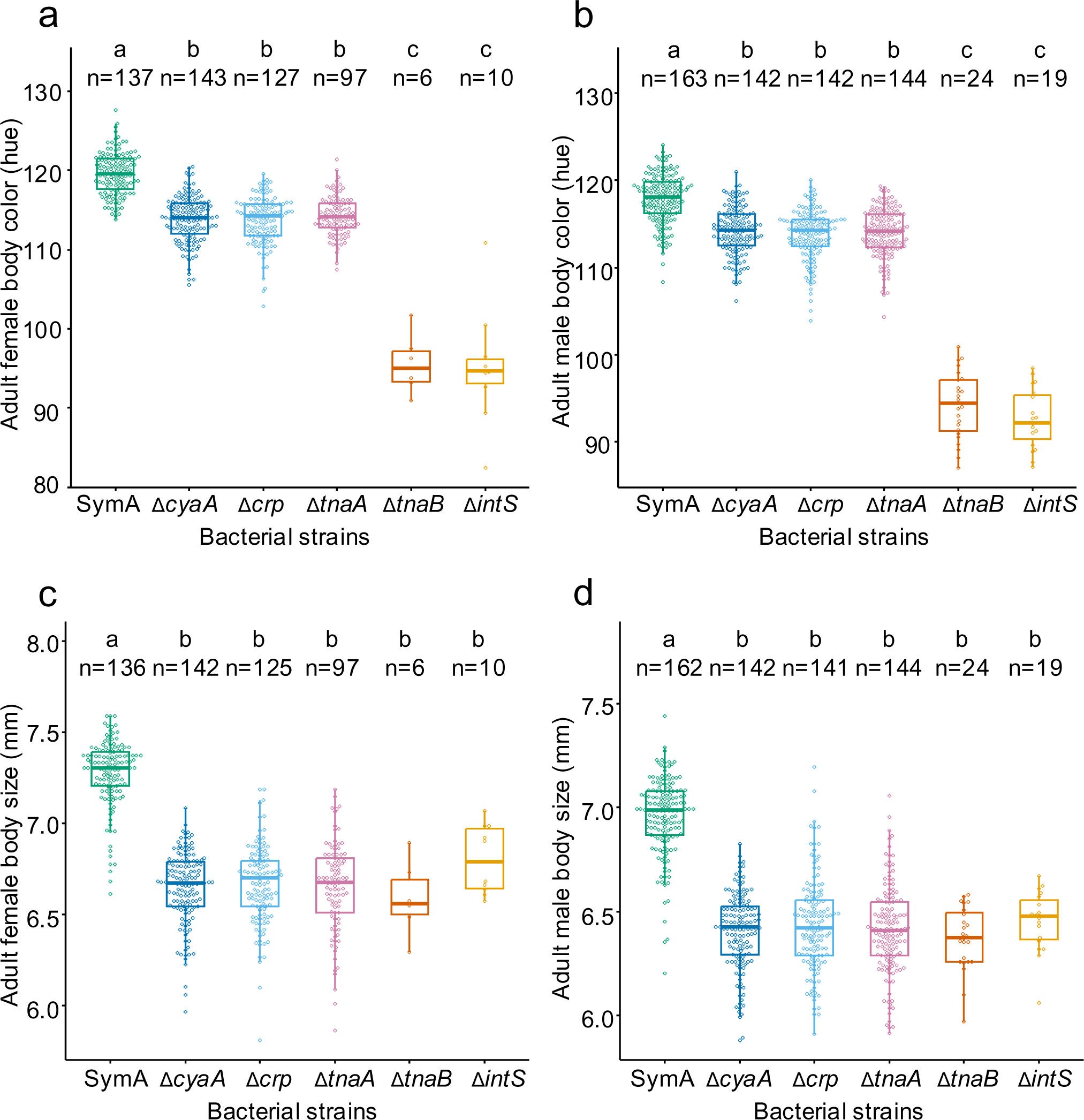
Phenotypic effects of *P. stali* infected with a natural symbiont *Pantoea* sp. A (Sym A), mutant *E. coli* strains (Δ*cyaA*, Δ*crp*, Δ*tnaA* and Δ*tnaB*), and a wild-type *E. coli* strain (Δ*intS*). (**a**) Adult female body color. (**b**) Adult male body color. (**c**) Adult female body size. (**d**) Adult male body size. Different alphabetical letters (a, b, c) indicate statistically significant differences (pairwise Wilcoxon rank-sum test with Hommel’s correction: *P* < 0.05).

**Fig. S6.**
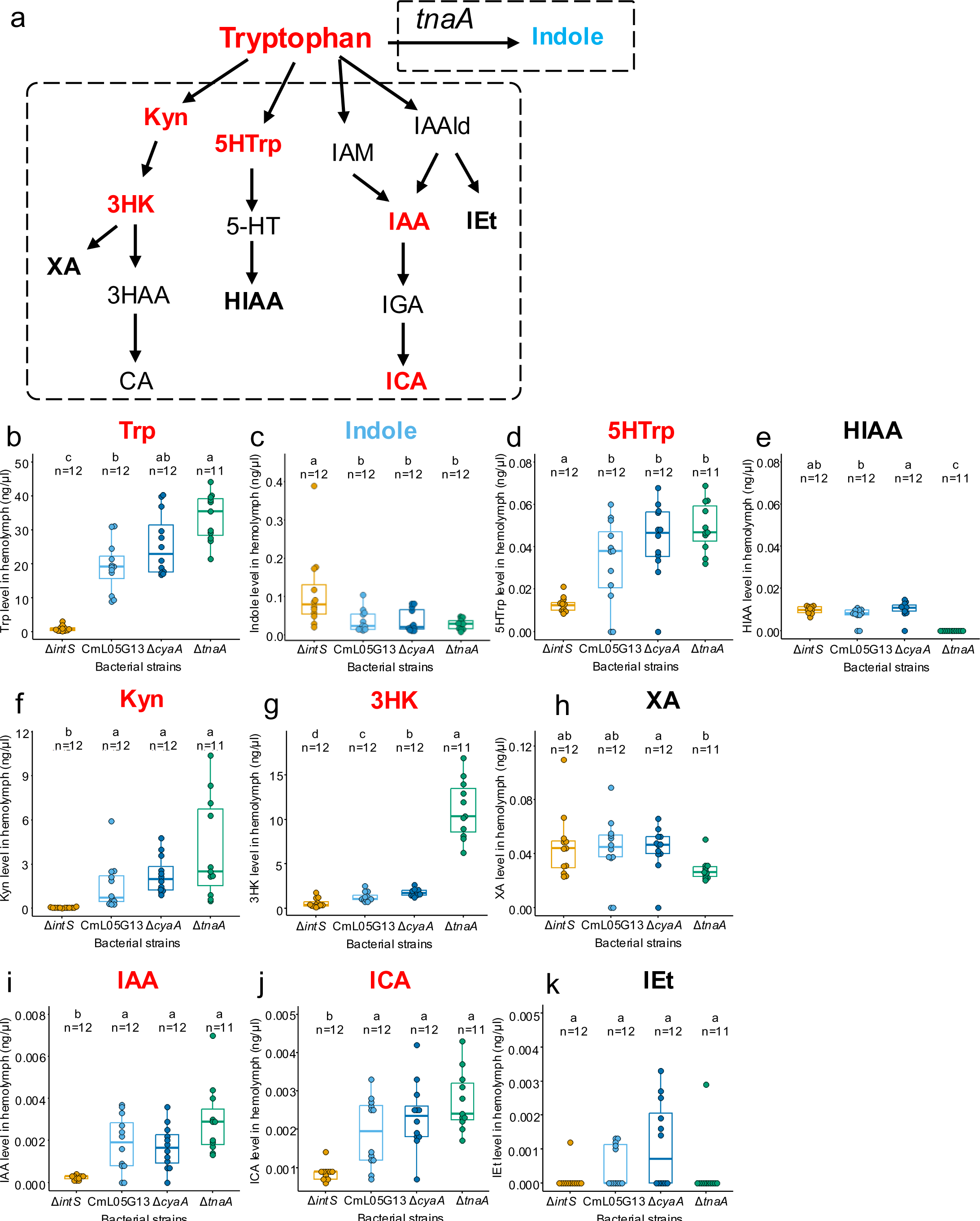
Titers of tryptophan-derived metabolites in hemolymph of *P. stali* infected with different *E. coli* strains. (**a**) A schematic overview of tryptophan metabolism. Compounds quantified by LC-MS are highlighted in bold. Red, blue and black represent metabolites that increased, decreased and unchanged by infection with the host performance-improving *E. coli* mutants (CmL05G13, Δ*cyaA* and Δ*tnaA*), respectively. (**b-k**) Titers of tryptophan and its derived metabolites in the host hemolymph. (**b**) Tryptophan (Trp). (**c**) Indole. (**d**) 5-Hydroxytryptophan (5HTrp). (**e**) Hydroxyindoleacetate (HIAA). (**f**) Kynurenine (Kyn). (**g**) 3-Hydroxykynurenine (3HK). (**h**) Xanthurenic acid (XA). (**i**) Indole-3-acetic acid (IAA). (**j**) Indole-3-carboxylic acid (ICA). (**k**) Indole-3-ethanol (IEt). Other abbreviations: 3HAA, 3-Hydroxyanthranilic acid; CA, Cinnabarinic acid; 5HT, 5-Hydroxytryptamine; IAM, Indole-3-acetamide; IAAId, Indole-3-acetaldehyde; IGA, indole-3-glyoxicacid. Different alphabetical letters (a, b, c) indicate statistically significant differences (pairwise Wilcoxon rank-sum test with Hommel’s correction: *P* < 0.05).

**Fig. S7.**
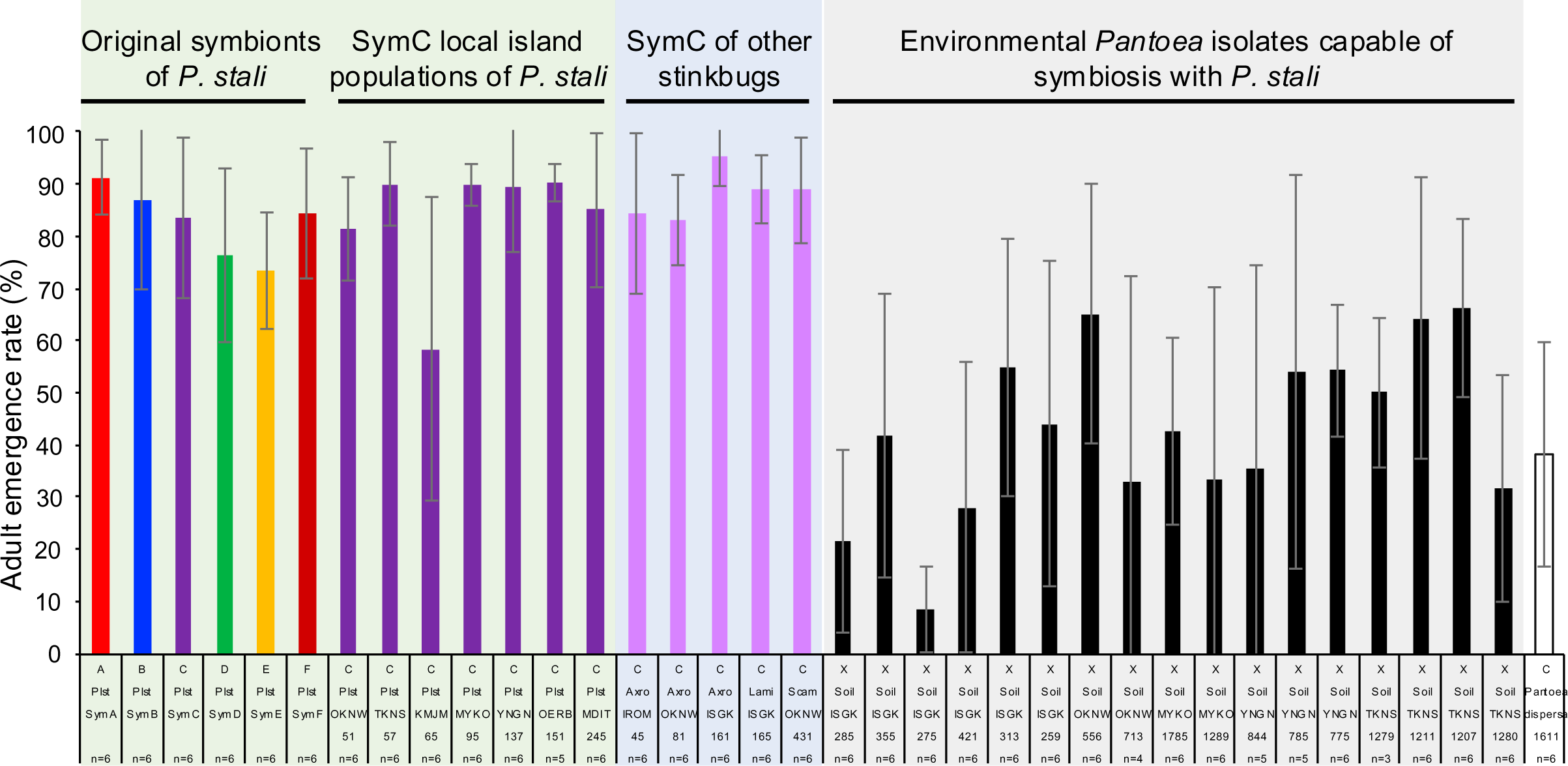
Adult emergence rates of *P. stali* infected with original *Pantoea* symbionts, heterospecific *Pantoea* symbionts from different stinkbug species, and potential *Pantoea* symbionts isolated from environmental soil samples.

**Fig. S8.**
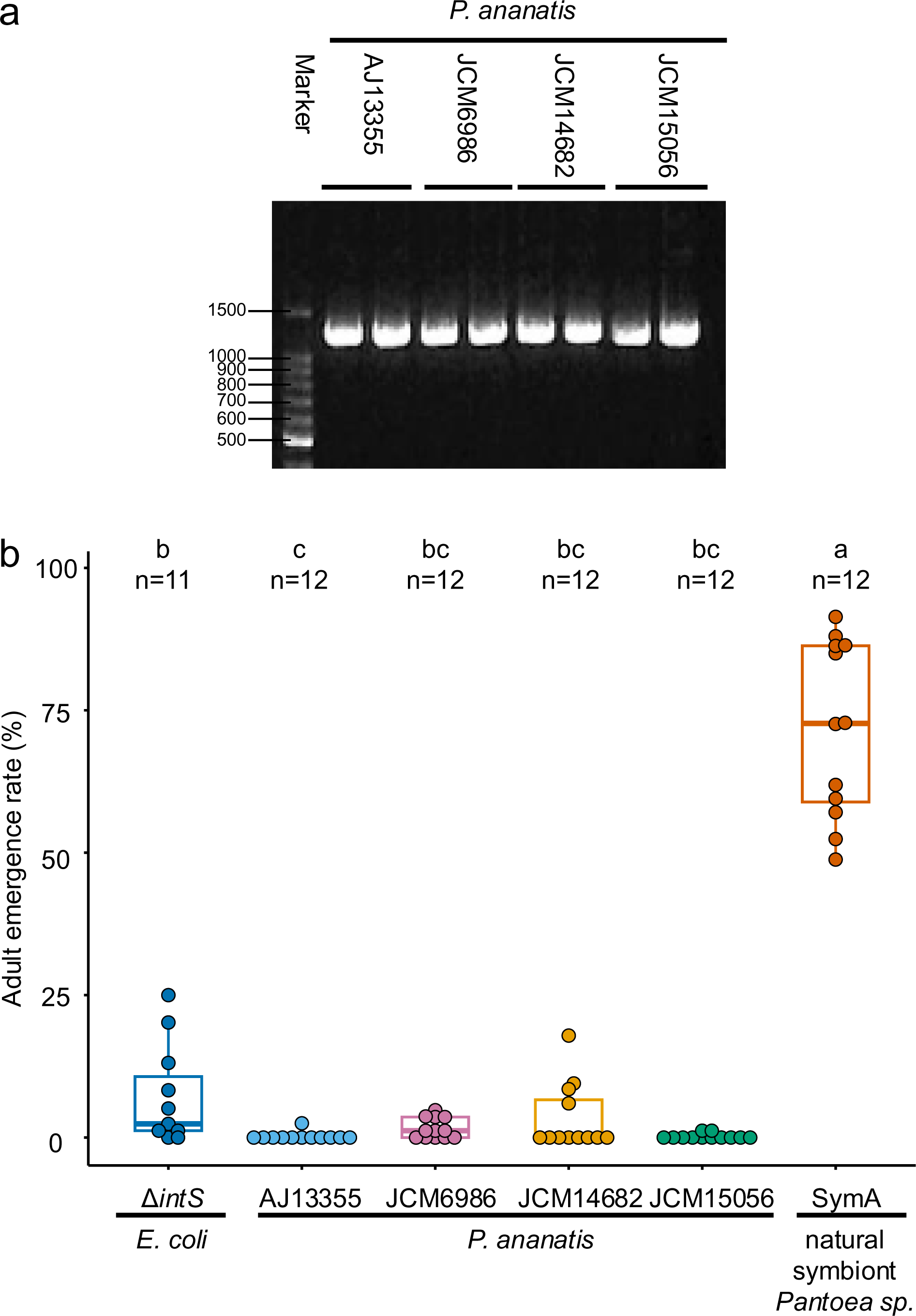
Presence of *tnaA* gene and symbiotic incapability of *P. ananatis* isolates. (**a**) PCR detection of *tnaA* gene in the *P. ananatis* isolates AJ13355, JCM6986, JCM14682 and JCM15056. (**b**) Adult emergence rates of *P. stali* infected with the *P. ananatis* isolates. The emergence rates were zero or near-zero for the *P. ananatis* isolates, which were drastically lower than the rate for the natural symbiont Sym A and even lower than the wild-type *E. coli* Δ*intS*. Different alphabetical letters (a, b, c) indicate statistically significant differences (pairwise Wilcoxon rank-sum test with Hommel’s correction: *P* > 0.05).

**Fig. S9.**
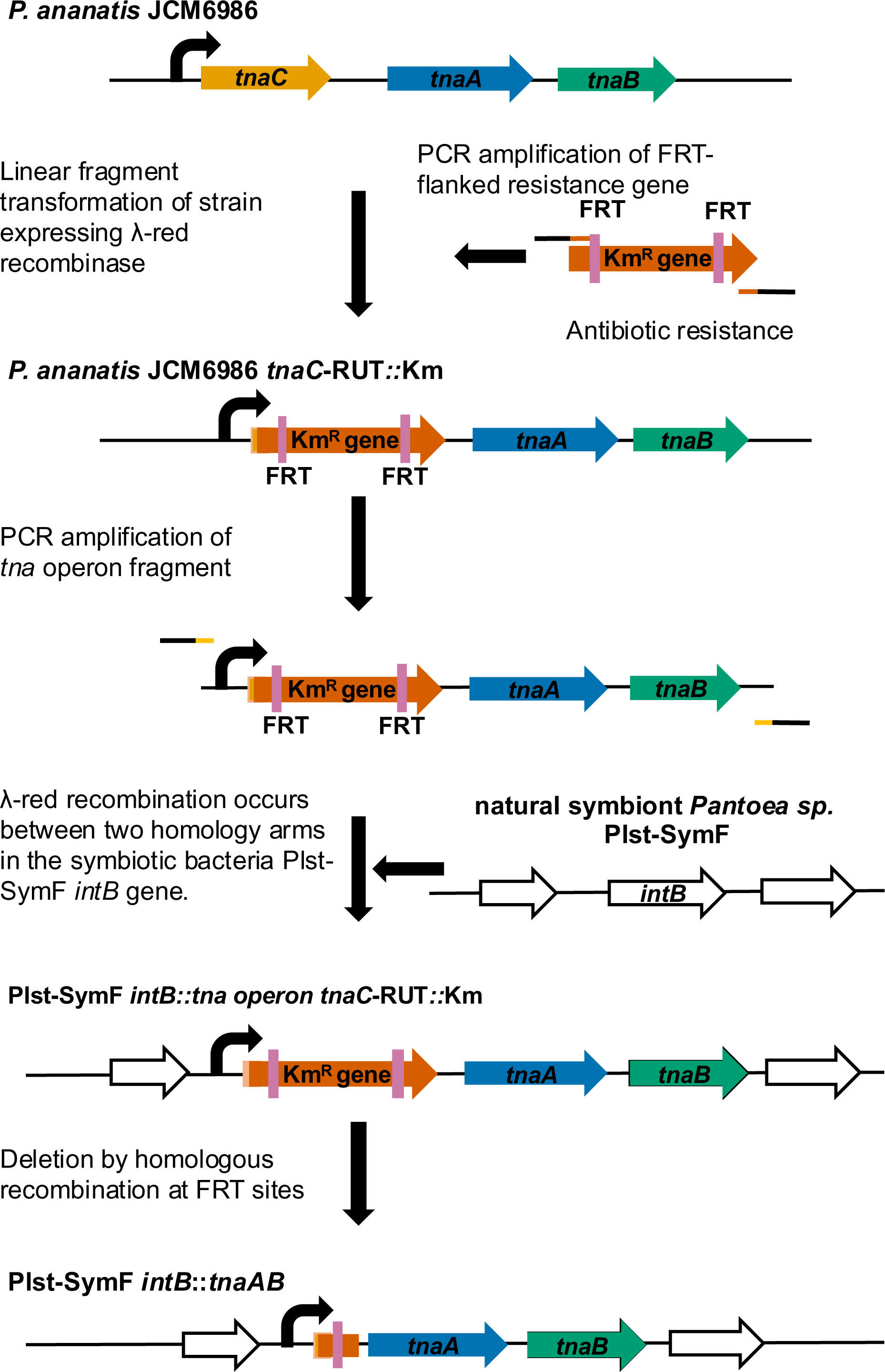
Schematic procedure for transformation of natural symbiont *Pantoea* sp. F (Sym F) with functional tna operon from P. ananatis.

**Fig. S10.**
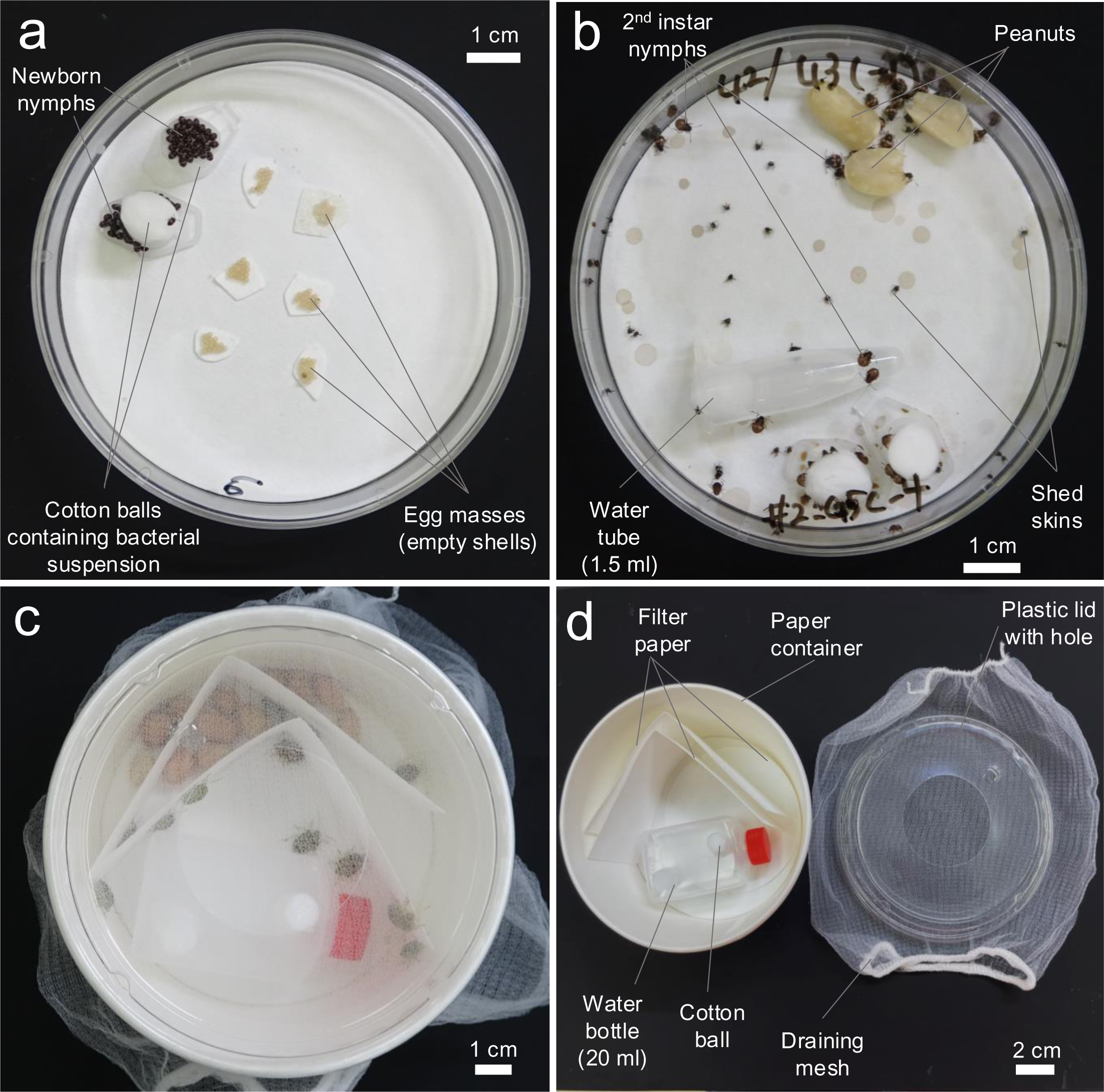
Rearing and inoculation systems for *P. stali*. (**a**) Petri dish rearing for newborn nymphs. Sterilized egg masses are placed in a clean Petri dish with two cotton balls on plastic tube lids. Upon egg hatching, 1.5 ml of bacterial suspension is added to the cotton balls, from which the newborn nymphs orally acquire the bacterial cells. (**b**) Petri dish rearing for early second instar nymphs. Upon molting to second instar, sterilized peanuts and a water tube are introduced, from which the insects start feeding. (**c, d**) Paper container rearing for late second instar to adult insects. After keeping in the Petri dish system for 3-4 days, the late second instar nymphs were transferred to a clean paper container with a water bottle and sterilized peanuts. Thereafter, the insects were maintained in this rearing system until adulthood, with renewing the container, water and food every week. The large paper container with a large-holed lid covered with mesh is for reducing the insect density and controlling the humidity, which is important for healthy survival of the insects.

